# The Swedish example of food animal production without extensive use of antibiotics – or “healthy animals do not need antibiotics”

**DOI:** 10.1101/809079

**Authors:** Ingeborg Björkman, Marta Röing, Jaran Eriksen, Cecilia Stålsby Lundborg

**Author notes:** **Correspondence:** Jaran Eriksen, Department of Public Health Sciences, Karolinska Institutet, Tomtebodavägen 18A, 171 77 Stockholm, Sweden.

## Abstract

**Objective:** To describe how stakeholders at different levels in food animal production in Sweden work to contain antibiotic resistance, with a special focus on poultry production. The stakeholders’ perceptions of antibiotic resistance and awareness of the One Health concept were also studied.

**Methods:** This is an interview study with thirteen informants. They represent policymakers, trade organisations, and veterinarians and farmers in the poultry industry. Interview transcripts were analysed using content analysis. The analysis continued until a latent theme emerged, and then the content was rearranged in four domains.

**Findings:** A latent theme “Working in unison” emerged, based on the consistency expressed by the informants when they discussed antibiotic resistance, use of antibiotics and food animal production methods. The theme was built on four domains, representing the content of the interviews: Knowledge and engagement; Cooperation; Animal health concept; and Development in balance with economic prerequisites. The work for healthy animals started in Sweden already in the 1920-ies and continued step by step in cooperation and with support from the government. In 1986 Sweden became the first country to ban antibiotics for growth promotion. Veterinarians were considered important drivers of processes by spreading knowledge and working close to the farmers. Farmers felt involved in the development of production methods. The One Health concept was well known among stakeholders working at national level but not among veterinarians in production or farmers.

**Conclusions:** Sweden has come far in work to contain antibiotic resistance in the animal sector by practicing restrictive use of antibiotics in food animal production. This practise is based on a long tradition of cooperation among stakeholders, from policymakers to farmers, and with a primary focus on animal health and welfare.

## Introduction

Antibiotic resistance is a growing global health problem threatening human and animal health (1,2) and was in 2013 ranked as the third worst global risk (3). In recent years new global risks, such as extreme weather events and failure of climate-change mitigation and adaptation, have emerged, and antibiotic resistance seems to be forgotten (4). However, antibiotic resistance is not slowing down according to a recent report from the European Centre for Disease Prevention and Control (ECDC) and the European Food Safety Authority (EFSA) (5) and efforts to contain antibiotic resistance are still urgent. In 2015 the World Health Organisation (WHO) announced a Global Action Plan based on a “One Health” approach (6). This approach was taken since resistant bacteria can be transmitted between humans, animals, food and the environment, and across international borders. This action plan emphasises a need for coordination among international sectors and actors including human and veterinary medicine, agriculture, environment, finance and consumers (6).

Efforts to contain antibiotic resistance started early in Sweden. In the human health sector “Strama”, the Swedish strategic programme against antibiotic resistance, was formed in 1995 (7). Even earlier, in 1986, the use of antibiotics for growth promotion in food producing animals was banned (8). Today Sweden has low levels of antibiotic use, and one of the lowest levels of antibiotic resistance compared to most countries in the world (9). The Swedish government strategy for containing antibiotic resistance from 2016 takes a One Health approach (10) with the overall goal to preserve the possibility of effective treatment of bacterial infections in both humans and animals. The strategy was up-dated in 2017 including an emphasis on international cooperation (11).

To be able to contain antibiotic resistance, knowledge and social engagement, as well action from different levels of society is needed. Although knowledge at research and policy levels has been available a long time, actions are still insufficient, and the problem of resistance is growing (2). It is therefore important to study how knowledge and action plans are transformed to practice.

This study is part of the ABRCARRO (A One Health Systems and Policy Approach to Antibiotic Resistance Containment: Coordination, Accountability, Resourcing, Regulation and Ownership) - an international project which aims to explore and describe how national action plans on antibiotic resistance were developed, implemented, monitored and evaluated in Sweden, South Africa and Swaziland. The project includes interviews with different categories of stakeholders, at government level, for example policymakers, and professionals in human, animal, and environment/agriculture sectors, as well as policy document analyses. The present study focuses on efforts to contain antibiotic resistance in the animal sector in Sweden.

## Method

This is an interview study exploring how activities to contain antibiotic resistance have been developed, implemented, monitored and evaluated in food producing animals in Sweden. A strategic sample of informants was recruited. The purpose was to gain a rich material of different perspectives and diversity. Informants were professionals at policy level, from authority or trade organisations, and practitioners. We chose to use poultry production as an example and practitioners were veterinarians in production and poultry farmers. A total of 13 persons were interviewed, see Table 1. All interviews were carried out by one of the authors (IB) between January and June 2018. The interviews lasted between 40 and 96 minutes, on average 62 minutes. Policy persons and persons at trade organizations were contacted via email, informed of the purpose of the study and asked to participate. Snowballing was used to find practitioners, both veterinarians in production and farmers. An additional informant, an egg farmer, was recruited via direct contact with a local farmer. Informed written consent was obtained from all. Ethical approval was applied for. According to an advisory opinion from the Regional Ethics Board in Stockholm, there were no ethical objections to the study (Reg number: 2017/1999-31).

**Table 1.** Description of informants.

| Level | Informants |
| --- | --- |
| Policy level<br>(3 informants) | Two veterinarians working at the National veterinary institute, one an expert on antibiotic resistance and one an expert on treatment of ruminants, and one veterinarian responsible for antibiotic resistance at the Swedish Board of Agriculture. |
| Trade organisations<br>(3 informants) | Head veterinarian at the Trade organisation for chicken production. Head veterinarian at the Trade organisation for egg production. Head veterinarian at The Federation of Swedish Farmers. |
| Practitioners in poultry<br>(3 + 4 informants) | Three veterinarians in production; breeder or hatchery. Three poultry farmers and one egg farmer. |

A semi structured interview guide was developed, the main questions are listed in Table 2. It was based on an interview guide previously used by the research group when studying perceptions of antibiotic use and antibiotic resistance. The questions were adjusted to focus on the One Health approach. The interview guide was pilot tested with two informants, one from the animal sector and one from human sector (human sector study is presented elsewhere). This pilot test did not change the interview guide and these informants were included in the studies.

**Table 2.**
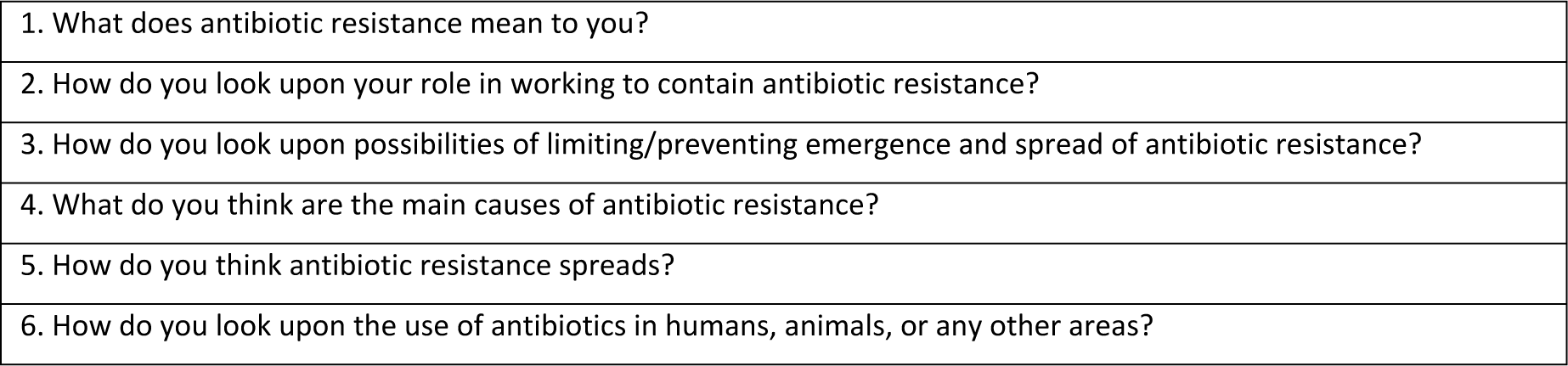
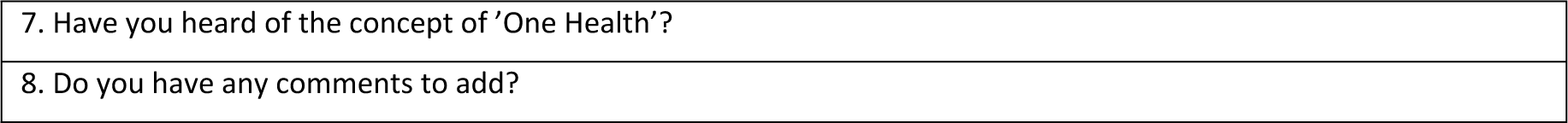
Interview guide used for interviews, main questions.

The interviews were performed at a place convenient for the informant, often at their workplace. The informants could associate and speak freely from the main questions, and the interviewer followed the conversation and asked probing questions. All interviews were audio-recorded and transcribed verbatim by an external transcriber. Before the analysis started, the interviews were listened through and transcripts checked by author IB.

### Inductive content analysis

One of the authors (IB) analysed the interviews. No theories or predefinitions were used, and conventional inductive content analysis was chosen (12). At start two of the authors (IB and MR) read the same transcript and marked meaning units and wrote preliminary codes. Then the researchers met, discussed and agreed on how to continue with the analyses. A first scheme of codes was constructed. Then IB proceeded and finished the analysis and the two researchers met several times and discussed the process and findings. Finally, all researchers discussed and agreed on the findings.

First the interview was read through to grasp the meaning. Then the texts were processed line by line and meaning units were picked from the text using the first scheme of codes previously constructed by authors IB and MR. The codes were used to sort the content from all interviews. In the next step the content of the codes was condensed, and a description was assigned to each code. The codes were grouped in comprehensive categories. Next the codes were condensed again, rearranged and merged. During this process a latent theme built on four domains emerged. In a final step, categories with their contents were rearranged in the four domains.

## Findings

The results of the content analysis are presented in Table 3, which shows the relation between theme, domains and categories, followed by a description of each domain and its categories. The descriptions in general represent the whole informant group, and focus primarily on poultry production. When a statement is related to a specific category of informants, the category is given, e.g. the category “veterinarian/s” is used when veterinarians from different categories express a similar perception. Descriptions are then followed by table 4, which presents quotes from the informants in the domains.

**Table 3.**
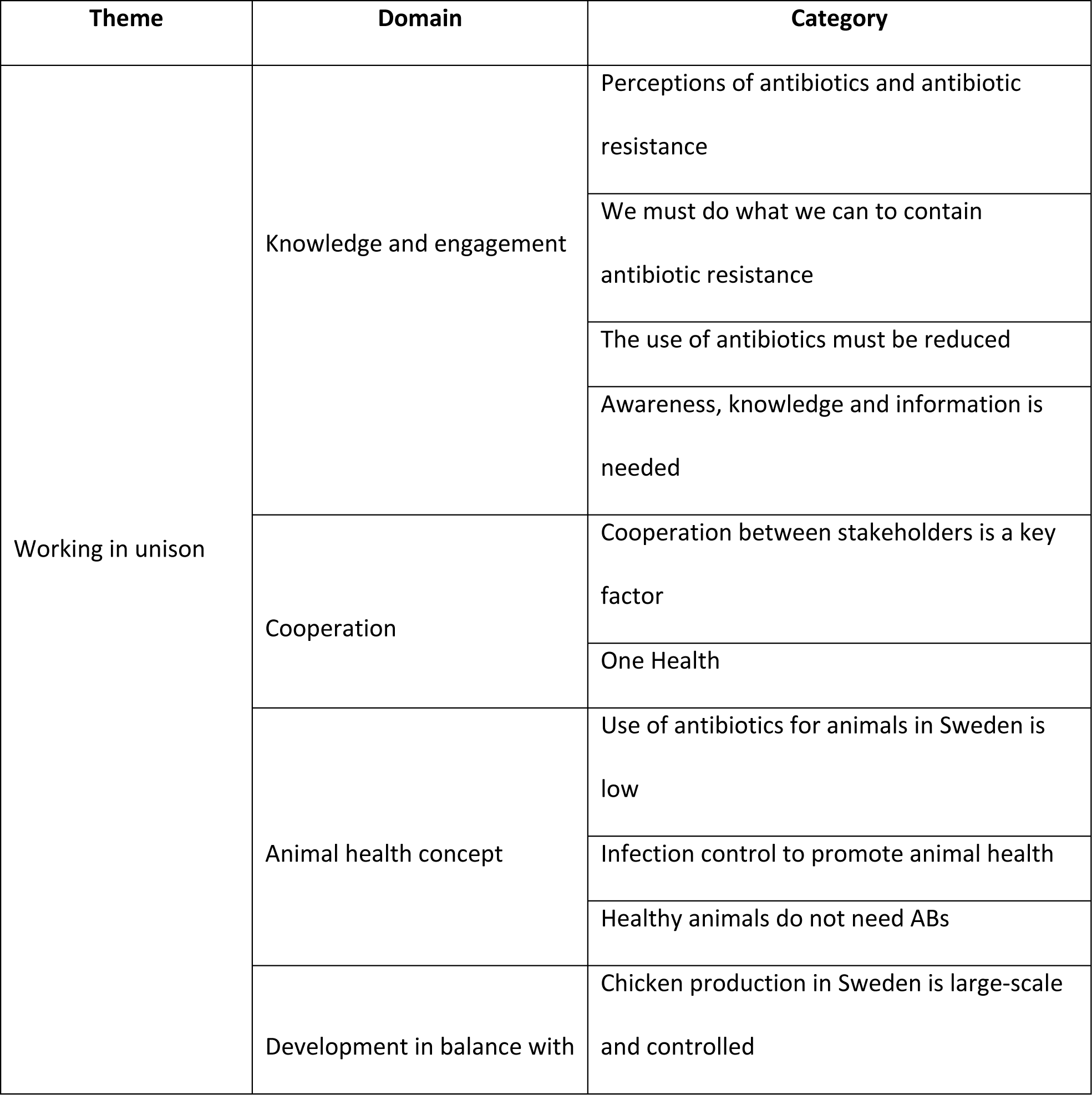
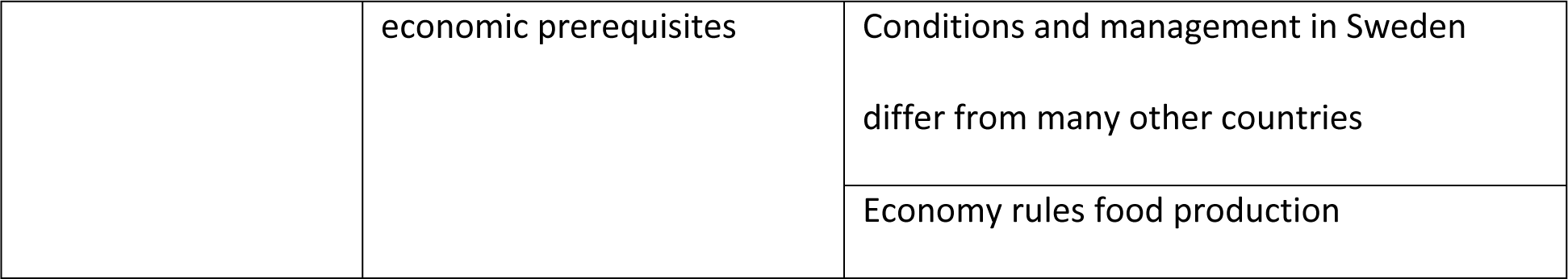
The categories are sorted in the domains that were identified together with the latent theme.

**Table 4.** Quotes from the informants sorted in domains. All the informants have contributed to the following quotes.

| Domain | Quotes |
| --- | --- |
| Knowledge and engagement | We have two possible tools we can work with - wise antibiotic use, and we can work with preventing infections. And then it is not only the spread of resistant bacteria, but all kinds of infections. [...] As it looks today, we can't **afford** not to work with both tools, and I don't believe, I don't believe it will be as effective if we don't work with both.<br>*Policymaker informant 1* |
|  | So we try, oh, oh...yes...to keep discussions alive during the whole year, both about disease control but above all the use of antibiotics in this area. *Trade organization informant 1* |
|  | When I write texts about this, I rarely need to change much if I publish it in the agricultural press or veterinary press. Because we both have great knowledge, and we are well aware of the issue.<br>*Policymaker informant 2* |
|  | But we try to work in such a way so that we don't use it [antibiotics], because - actually we think somehow that...it is not necessary [in chicken production] - it is instead very much about management factors. *Veterinarian in production 3* |
|  | So all the breeders really work to minimize the risk of contamination in the stable. So we change clothes completely and yes, or... you have done something so you wash your hands once again if you want, but now it is a fairly clean environment here so to speak, and then shoes are changed once more as well. *Farmer 2* |
| Cooperation | Everyone, everyone owns it [the antibiotic resistance issue]. And that's what I think we are so successful with in Sweden, eh, that we ... If I look at the animal sector, then it is really that we veterinarians work **together** with the farmers a lot in this matter. *Policymaker informant 2* |
|  | But I can never communicate, succeed in communicating with all Swedish veterinarians and farmers. Possibly with veterinarians, but not farmers. And they are the ones we need to reach in the end. Eh...they also need knowledge. And then one must work, must and must, but then my idea is to work via contacts, which is most effective, and to do so in close agreement with them.<br>*Policymaker informant 1* |
|  | In our field we have been quite skilled at cooperating with authorities, I think, and have developed a lot of these different programs to ensure the quality we have. *Farmer 1* |
|  | Facilitating, it's, that we are so ... have so much in common and cooperate, so that everyone doesn't need to do it at home in their house, but that we can actually share, so if the other company does tests to see if you can hatch chicken without this bacterium, for example, just by a very fine egg quality, they share the result so that we others can see it. *Veterinarian in production 1* |
|  | You have to work together, eh, so that you, as a rich western country, do not just sit on your high horse and eh, judge and point with your whole hand and say that now you should do this. On the contrary, you have to actually help. *Trade organisation informant 2* |
| Animal health concept | It is very rare that I have used antibiotics during the time I have done this [chicken-meat production].<br>Interviewer: Mm, how long is that?<br>Yes, it is almost 20 years, I think I can count on one hand what I have been prescribed [to the animals]. *Farmer 3* |
|  | There is a constant struggle with maintaining biosecurity, mentality, education, new people keep coming, experienced people quit, and you have to keep the flow moving. *Veterinarian in production 2* |
|  | Sweden as the first country banned antibiotics as growth promoters in 1986. And then you **saw**, it wasn't just... eh, the effects were quite big, because it was not just growth promoting, that antibiotics smoothed **over** management flaws. *Trade organisation informant 3* |
|  | There, like, the state did not go in, but the industry decided, it was an agreement then, that farmers, veterinarians, veterinary organizations decided that we should, we should eradicate it [Bovine virus] from the country. And then it was a voluntary control program.<br>*Policymaker informant 2* |
| Development in balance with economic prerequisites | We have a salmonella control program for example, you can eat your eggs raw in Sweden, you can do quite a lot that you cannot do in other countries. And it is something that has cost money, so this has been yes, it has been done with government grants.<br>*Policymaker informant 3* |
|  | The risk is, if you go too fast [in changing production methods], you get setbacks and then the producer says that this is not possible, it's not possible - and then you are back where you started.<br>*Trade organisation informant 1* |
|  | Yes, I think we have really good breeders, most of them. And they, they want, for their own sake, it is very much about avoiding salmonella, after all, and it goes hand in hand as well ... Salmonella and campylobacter for them are the ones they work, they get deductions if |
|  | they have campylobacter, and [if they have] salmonella so the whole herd is slaughtered. *Veterinarian in production 3* |
|  | Of course it is important that we are compensated for the extra cost eh, that this system in this case has cost us, and partly it is about communication with the consumer and explaining why this is a little bit more expensive yet has these advantages, and one may well need help from authorities and politicians as well as to explain and describe. *Farmer 1* |

### 1) Knowledge and engagement

#### Perceptions of antibiotics and antibiotic resistance

All informants, except one of the farmers, were engaged in the question of antibiotics and antibiotic resistance. All the other twelve informants shared the perception that antibiotics are needed but must be used restrictively. Farmers mainly said antibiotics are necessary for humans, while veterinaries said for both human and animals. A few acknowledged the risk of underuse of antibiotics in humans and animals. Veterinarians stressed that if animals are ill and treatment is available, antibiotic treatment must be given. Both veterinarians and farmers emphasized the importance of developing new antibiotics in need of more resources, and that this is a political issue.

Except for one of the farmers, the informants were very concerned about antibiotic resistance, and described it as a very serious threat. They understood that antibiotic resistance means that we cannot treat diseases, not perform surgery safely, as well as increased mortality. A common perception was that antibiotic resistance already exists, but that the real threat is a future problem. One policymaker informant pointed out the economic consequences of antibiotic resistance and referred to estimates from the World Bank Group. A farmer thought that soon people will hesitate to travel abroad, due to the risk of bringing back antibiotic resistance. Some of the veterinarians compared the antibiotic resistance issue with the issues of environment and climate – creeping threats, and issues which bring out the need for behaviour change in humans. Furthermore, climate change and antibiotic resistance were said to be connected, and climate change can increase the problem of antibiotic resistance.

Informants perceived antibiotic resistance as caused by too extensive consumption of antibiotics in the human sector, but the animal sector also contributes to the development of antibiotic resistance. They also felt that antibiotic resistance mainly developed abroad and then imported to Sweden. Veterinarians had comprehensive knowledge and could explain how antibiotic resistance develops. Some veterinarians explained that resistant bacteria we bring home when travelling disappear after a while. One policymaker informant asked for more knowledge about how chemicals as heavy metals and biocides can stress bacteria into developing resistance.

#### We must do everything possible to contain antibiotic resistance

All veterinarians emphasized that we must do what we can to contain antibiotic resistance. In their opinion, this meant to reduce antibiotic usage and use antibiotics wisely. This was everyday ongoing work, expressed one policymaker informant, and the work must be done in both animal and human sectors. All informants perceived antibiotic resistance as being an issue for everyone – everyone must engage and authorities, trade organisations, as well as farmers must be involved. Physicians and veterinarians must take responsibility for not prescribing antibiotics unnecessarily. Treatment of pets was a special issue according to some veterinarians, as animal owners may demand antibiotics for their pets. Veterinarians need to agree on being more restrictive, and at the same time, acknowledge that pets nowadays are perceived as family members. Veterinary competence was present in authorities, trade organisations and in the production chain. A future possible threat was lack of competent veterinarians, as young veterinarians prefer to work with pets and horses instead of farm animals.

#### The use of ABs must be reduced

The veterinarians explained that eradicating antibiotic resistance is difficult but that reducing antibiotic use is possible. The purpose is to reduce selection pressure. The use or non-use of antibiotics in food producing animals is a matter of production methods, veterinarians said.

Methods to reduce antibiotics in food producing animals were similar to methods in humans, e.g. finding alternatives to antibiotics, refraining from antibiotics when treatment is not necessary, and practicing good hygiene, prevention and infection control. Veterinarians, before choosing antibiotic treatment, always tested for antibiotic resistance. One veterinarian used ‘wait-and-see’ instead of prescribing antibiotics and offered a second visit at the farm some days later to check up on the animals. Narrow-spectrum antibiotics were used if treatment was deemed necessary. Some veterinarians worried that narrow spectrum antibiotics would be removed from the market, because of limited use in some countries and a small production.

Monitoring was an important part of antibiotic resistance work, the veterinarians said. Most important was to follow the use of antibiotics, which shows where efforts to reduce antibiotic use are needed and demonstrates possible effects of efforts. Since year 2010 the trade organisation “Svensk Fågel” (translates to Swedish Bird/poultry) collects statistics on antibiotic use in poultry. Veterinarians at trade organisations wished for more developed statistics on antibiotic use for benchmarking.

#### Awareness, knowledge and information is needed

The informants expressed that awareness, knowledge and understanding was necessary among stakeholders and the public to make people follow available recommendations. The perception was that Swedes in general were aware of antibiotic resistance and that this facilitates work to reduce antibiotic use. Veterinarians thought media can contribute to awareness of antibiotic resistance among the public and drive work to reduce antibiotics forward. This was confirmed by the farmers, who referred to media when they described what they knew about antibiotic resistance. However, it may be difficult to understand the real significance of the threat of antibiotic resistance, several informants pointed out. The message was difficult to communicate, one policymaker informant said. The informants gave examples of outbreaks of infections that had increased awareness, both in Sweden and internationally, which had led to measures to reduce the risk of spreading infection. The informants believed awareness internationally in general was lower than in Sweden.

### 2) Cooperation

#### Cooperation is a key factor

An important experience reported was the close cooperation that existed among authority, academia, trade organisations and other stakeholders in the animal sector in Sweden. Veterinarians mentioned this as a facilitating factor. Even farmers felt included and pointed out that the different actors in the poultry industry had developed the production methods in cooperation with each other. One farmer explained that regulations set up by authorities without cooperating with chicken farmers would not work, since farmers need to have the same picture, to understand the whole. The trade organisation organized all actors in the production chain, forage producers, farmers, veterinarians in production, and slaughterhouses. Veterinarians in poultry were few, they met at the trade organisation and agreed on how to act. Some veterinarians in production said their company directors may compete, but when it comes to veterinary medicine the veterinarians cooperate. Chicken farmers were also rather few and met regularly at the trade organisation’s yearly training days. Knowledge on good production methods was easy to spread.

Cooperation in Sweden between animal and human sectors at policy level has a longstanding history, in Strama since the 90s, and later at the Swedish cooperative platform. Here animal and human sectors agree to reduce the antibiotic use. A rather new discussion was cost sharing, meaning that costs were to be shared by both sectors when actions were taken in the animal sector for the sake of public health.

One hindering factor expressed by the veterinarians was the ‘blame game’. This meant blaming other professionals, sectors or other countries for doing less, a belief that others must change but that we are doing enough. This could happen between animal and human sectors, both locally and internationally, or when statistics on antibiotic use were presented and countries were compared. Such attitudes could hinder the will to cooperate and stop efforts to reduce antibiotic use.

#### One Health

Sweden has established a cooperative platform, commissioned by the Swedish government, which has reduced the blame game between sectors, according to one policymaker informant. Recently new sectors have joined in, as directed by the government. The platform works according to WHO’s Global Action Plan and the One Health concept. Policymaker informants spontaneously mentioned they had a role in One Health. The concept was well known among trade organisation informants. Two veterinarians in production had heard about the concept but were not exactly sure about the meaning. None of the farmers had heard of the concept.

Two policymaker informants knew that antibiotic resistant bacteria can be found in the environment, and that the meaning of this is not yet known. They explained that we need more knowledge to understand the meaning and how the environmental sector shall be involved in the One Health work. The other informants had no knowledge of antibiotic resistance in the environment but reflected on the issue.

### 3) Long tradition of animal health concept in Sweden

#### Use of antibiotics to animals in Sweden is low

All informants talked about how little antibiotics are used in food producing animals in Sweden. Antibiotic use was especially low in chicken farming and was not used at all in egg production. Areas for improvement in animal sector were mentioned, including antibiotic treatment of pets, the use of coccidiostats in chicken, and veterinarians trained outside Sweden who often had other views on antibiotic treatment and prescribed antibiotics more often than Swedish trained veterinarians.

The informants did not see antibiotic resistance as a problem in food animal production in Sweden. The farmers said it did not affect their work. However, all informants talked about the fact that in 2010 Swedish chickens were infected by ESBL from breeding animals. Despite hard work Swedish chickens may still carry ESBL. As one veterinarian in production explained, *E coli* infections in chickens are not treated with antibiotics so as not to promote spread of ESBL resistance.

All informants perceived antibiotics to be extensively used in food producing animals abroad. One veterinarian reported that 90 percent of all antibiotics for animals in Europe were used for herd treatment. However, it was also pointed out by veterinarians that now more countries are working hard to change their food production and use less antibiotics.

#### Infection control to promote animal health

All informants talked about the importance of preventing spread of infection. Veterinarians emphasized that this was a way to reduce the need for antibiotics and argued it was economical to prevent infections. One policymaker informant commented that at a global level, sanitation improvement and hygiene in humans and good manure management would reduce the risk of spreading diseases in both animals and humans.

Veterinarians described measures to prevent spread of infections, e.g. contact isolation, to put down animals, to limit trade to regions that are not infected, and to practice biosecurity. Veterinarians in production and farmers gave detailed descriptions of how they worked with biosecurity. They said that biosecurity was well established and followed by all poultry farmers. Biosecurity had been in use since the 80s, “this is how we work, all farmers in poultry do it”, one farmer expressed. One veterinarian in the production described the requirement to hire a person responsible for infection control, and to establish good routines for hygiene, keep records, and to follow standards for the stables, including hygiene barriers and visit restrictions. Veterinarians had continuous training for new staff at the farms.

Veterinarians explained that Sweden has eradicated several diseases in farm animals. The first was tuberculosis in cows in 1920s, a governmental initiative. Another example was bovine virus in cattle, for which a voluntary infection control program was developed by farmers, veterinarians and veterinary organizations. When vaccines are not available, put down animals have been used, which is also a method to containing susceptibility to penicillin. One veterinarian at a trade organization described how tough this work was for farmers. Sometimes, a farmer’s whole livestock was put down, and years of breeding work destroyed.

One trade organization informant mentioned that to protect Swedish animals from infections, legislation restricts import of animals and breeding material. Furthermore, voluntary import restrictions have been added by the trade organizations. Since 1995 Sweden belongs to the EU market but has so far been allowed to keep this stricter regulation, thanks to the successful eradication of diseases. In 2013 Sweden was the first country to launch a legislation of infection control in veterinary medicine. This legislation was prepared by one of the policymaker informants.

#### Healthy animals do not need antibiotics

A facilitating factor in the efforts to contain antibiotic resistance, according to one policymaker informant, was that antibiotic resistance has never been looked upon as an isolated issue but as part of a whole, a bigger picture. All informants pointed out that animals who are well cared for feel better and stay healthy. “Healthy animals do not need antibiotics” was repeated by many informants as a motto. Veterinarians claimed that healthy animals have strong immune protection and are more resistant to disease. To enable this, the farmers would buy chickens of high quality, use feed of high quality, impose careful infection control and keep good hygiene. Furthermore, they would change conditions if anything stressed the animals, e.g. the temperature in the stall, nutrition supply, water supply. Veterinarians and farmers stated keeping animals’ health was a daily never-ending process.

### 4) Development in balance with economic prerequisites for stakeholders in Sweden

#### Chicken production in Sweden is large-scale and controlled

Chicken production in Sweden was described by the informants as industrial, large-scale and well controlled. Globally, the chicken industry was described as a pyramid, with a few breeding companies on top producing grandparents to all chickens in the world. Sweden buys from two of these breeders. There are two levels in Sweden above the chicken farmers, breeders and hatcheries. The hatcheries deliver chickens to the farmers, which in turn are connected to a slaughterhouse. The slaughterhouse does a planning based on peoples’ demand of chicken and calculates the number of chickens to be ordered from the hatchery, and when they need to be delivered. After delivery the chickens live indoors until slaughter. Biosecurity was prioritized, and locally produced food was recommended before ecologic production, which was regarded riskier for chickens and too costly for many consumers. This was an overall perception among the informants, except one farmer who had small-scale ecological egg production.

Veterinarians talked about the long tradition of infection control programs in Sweden. An important step was taken after the Swedish so called ‘Alvesta epidemic’ in the 1950s when 90 people died due to salmonella. This outbreak started the development of a salmonella control program and animal welfare programs. A next important step was an initiative, taken by farmers, which led to the ban of growth promotion antibiotics in 1986. A consequence of not using antibiotics for growth promotion was the need to change production methods. It has been costly but now they see the benefits, said one trade organisation informant. One lesson in Sweden according to another trade organisation informant was that change must be allowed to take time. If progress is too fast, there can be rebound effects and producers may stop believing animal production without antibiotics is possible and may not want to cooperate. Many informants wished Sweden could be a role model for other countries and show that it is possible to change the food animal production.

#### Conditions and management in Sweden differ from many other countries

Informants said production methods for Swedish chickens differ from many other countries. By ‘other countries’ they usually meant the rest of Europe except the Nordic countries, but sometimes it included the rest of the world. As an example, it was stated that the maximum kilogram living chickens allowed per square meter was higher in other countries compared to Sweden. All countries in the EU have a common animal law but despite this, production methods and level of antibiotic use varies. A policymaker informant concluded that laws are obviously not enough to have an impact on the food animal production, there seem to be other factors that rule.

Both veterinarians and farmers talked much about their efforts to make animals feel good and be as healthy and strong as possible by focussing on prevention, biosecurity and animal welfare instead of using antibiotics. Veterinarians said Sweden benefited from having a cooler climate and seasonal variation, whereas the risk of spreading disease is higher in warmer countries. A facilitating factor was the protection of animal health by restrictive import. Bacteria in animal production is both a question of animal protection and of food security. If salmonella is detected the whole chicken herd is put down, but chickens with campylobacter are taken to slaughter. Bacteria die if meat is heated to a high enough temperature, and consumers need to know this.

#### Economy rules food production in Sweden

Economy rules the chicken production and the production methods. A farmer explained the economic interest to follow all control programs very carefully, especially when you have a large production it will be very costly if something goes wrong. A veterinarian in production reflected that it is not laws that rule production methods, it is profitability. One veterinarian believed that in Sweden, agreements and guidelines had been more important than legislation, while another believed that governmental financing had controlled the development of food production methods and there had been both legislation and voluntary actions. The government has contributed financially to eradication of diseases.

All informants felt that farmers must be able to live on their production. If they cannot sell their goods, production will cease. Buying Swedish meat supports a production that uses less antibiotics, and not buying Swedish products means moving the antibiotic resistance problem somewhere else. However, production without antibiotics was said to be more expensive and informants suggested that the Swedish government could support Swedish production by explaining to consumers why Swedish meat is more expensive. There was a belief that many Swedish consumers trust the Swedish production of meat.

Veterinarians expressed that we must safeguard the production we have, and governmental politicians must know this. Threats to the Swedish production were highlighted, for instance too strict regulations, lack of understanding of the factors that can undermine the food industry, and Swedish animal-rights organisations which work hard to eliminate Swedish animal food production.

Politicians have a responsibility for the antibiotic resistance issue. Regarding this, policymaker informants seemed to think globally and the other informants nationally or at EU level. Politicians need to allocate resources for research, monitoring and education. One farmer was sceptical and felt that politicians think too short-term. Containing antibiotic resistance is politically charged, one trade organisation informant said, reflecting on selling meat in the common market.

The informants thought it was difficult to assess whether enough resources were allocated to containing antibiotic resistance. Some policymaker informants mentioned that authorities needed more resources for their work. Informants from all categories acknowledged the cost of work to contain antibiotic resistance. One policymaker informant concluded that a country with less resources does not come as far, even if they would like to. Conditions differ and countries may have to prioritize other measures. There is a lot of money in the food industry, people have short-term thoughts of economic profit and do not believe that food production without antibiotics is possible. In addition, in certain countries veterinarians must sell medications to improve their salary and furthermore consumers want to buy low-cost meat.

Veterinarians expressed that Sweden and EU have an international responsibility to support containment of antibiotic resistance with funding and competence. Work in Sweden must continue, but we must also help other countries. Many informants called for research on which measures are most cost-effective.

## Discussion

This study gives insights into how stakeholders at different levels of the food producing animal sector in Sweden have been working together to develop production without extensive use of antibiotics. The measures taken have been successful. This seems to be due to a long-standing culture of cooperation between different stakeholders in Sweden. The latent theme “Working in unison” was based on the consistency expressed among the informants when they discussed antibiotic resistance, use of antibiotics and production methods, with a special focus on poultry.

The WHO guidelines for antibiotic use in food-producing animals include complete restriction of antibiotic use for growth promotion and disease prevention in healthy animals, and restrictive use of antibiotics identified as critically important for humans (13). Recommendations are based on evidence of decreased presence of bacterial antibiotic resistance in animals, and also humans, after interventions to reduce antibiotic usage (14). According to our findings, the WHO guidelines are followed in Sweden. Studies have shown that stakeholders in food production may believe they use less antibiotics than others (15). This could also be the case here. However, statistics on antibiotic use in food producing animals show that antibiotic usage in Sweden is low, only Norway and Island use less antibiotics (9).

### Key players – veterinarians and farmers

Regulations and action plans at global and national levels recommend restrictive antibiotic use in order to contain antibiotic resistance. To make a change, theory needs to be transformed into practice, and actors need to believe in the message. Some actors contest the link between agricultural antibiotic use and antibiotic resistance, but studies report compelling supporting scientific evidence for the need to take action (2,13,16). Another prevailing opinion is that the risk of developing antibiotic resistance is due to residues of antibiotics in meat. Even the meat industry has presented this perception, and argues that consumers do not need to worry because there are regulations on washout-periods after antibiotic use to prevent this happening (17). Although washout-periods do reduce antibiotic content in meat, this is not the whole issue of how antibiotic resistance is developed and spread. Residues of antibiotics have been detected in food in countries where regulations on antibiotic use are in there initial phases, i.e. India (18)

In chicken-meat production the suggested methods to decrease the need of antibiotics include biosecurity, hygiene, management, vaccine and probiotics (19,20). Farmers and veterinarians have been identified as key players in work to contain antibiotic resistance in the animal sector (15), and these were the stakeholders included in our study. What we found was veterinarians with high level of awareness of the threat posed by antibiotic resistance and in-depth knowledge of emergence and spread of antibiotic resistance. This was combined with a commitment to protect antibiotics while also protecting the animals and the food production. Furthermore, the veterinarians held positions at different levels where their knowledge and engagement came into use. A review including stakeholders primarily from countries in Europe and the US in pig, cattle and dairy farming concluded that veterinarians in general supported the reduction of antibiotics in food producing animals (15). However, some veterinarians believed that antibiotics for prophylaxis was judicious, and feelings of pressure from farmers, feed suppliers and others to use antibiotics were reported (15). An Indian study focussing on veterinarians in dairy farming, showed that veterinarians mostly prescribed antibiotics according to treatment guidelines, but also that they lacked restrictive antibiotic practices (21). A UK study used vignettes to explore which factors influenced veterinarians in their decision-making process. Several factors had significant influence, and included case type, farmer relationship, other veterinarians in practice, time pressure, habit, willingness to pay and confidence in the farmer (22). Motivation of misuse of antibiotics was studied by applying TPB (the Theory of Planned Behaviour) and found that US veterinarians were influenced by expectations from and obligations to different actors, i.e. other veterinarians, clients, consumers, pharmaceutical companies, and regulatory bodies (23). Hence, it seems important to reach agreement among the various stakeholders on how to produce animal products without extensive use of antibiotics, in order to help veterinarians prescribe antibiotics restrictively.

The farmers in our study said that they did not need to reduce the use of antibiotics, it was already zero or close to zero treatment. They were much more engaged in describing their daily efforts to keep the animals as healthy as possible. Farmers in other countries in Europe and the US acknowledge the need for antibiotic reduction, but some believe in the necessity of antibiotic use for a good profit in food animal production (15,24). To change farmers’ perceptions and practices, it has been suggested that veterinarians could play a role as sources of information and to facilitate learning processes (15,24). Networks of veterinarians and farmers may support such learning, and veterinarians need to understand this important function they have. Veterinarians in India working in dairy farming did not show awareness of establishing client relationship with stakeholders (21). In our study the close network of farmers and veterinarians was obviously important. The farmers often referred to local veterinarians and they also expressed trust in recommendations from authorities. Veterinarians at trade organisations seemed to play a role as a link between authorities and the farmers. Our study adds one factor, the farmers’ expressions of feeling involved in the development of production methods. This suggests that farmers were not only passive receivers of guidelines. On the contrary there seems to have been an exchange of information among stakeholders. Farmers were educated to understand the background of new management methods and they were given the opportunity to contribute with their knowledge from the field. Implementation research show that passive distribution of guidelines are ineffective and active measures are more successful (25,26).

### Consumers perceived both as a threat and as a possibility

With an increasing global population and subsequent increased demand for food, poultry farming has provided meat at a low cost in high-density poultry farms (19). However, a problem is that chickens often grow up in overcrowded stables, with poor hygiene and high risk of bacterial infections, and low doses of antibiotics are routinely given to prevent infections (19,27). As an example, a study on antibiotic use at eight chicken-farms in Thailand concluded that probably several tonnes of antibiotics are used every year in Thai poultry farming (28).

Products must be sold and thus, consumers’ buying choices have impact on how food is produced, and on how antibiotics are used in animal-food production (15,23). What is important for consumers? Price is mentioned routinely as a major influence. A study on consumers’ willingness when purchasing foods for a ‘sustainable diet’ identified high prices of recommended foods as a key barrier to change (29) and price was the highest limiting factor for buying organic chicken-meat (30). Another factor of influence is country of production. Some studies show that consumers appreciate domestically produced food, i.e. preferences for indigenous chicken meat and egg was high among consumers in Kenya (31) and consumers in Finland preferred Finnish produced broilers (32). However, exploring consumers from five different countries, Germany, France, Denmark, China and Thailand, and their choice of imported foods revealed that all these consumers tended to prefer foods from economically developed rather than less developed countries (33). The demand for organically produced food has increased in the last decades. This is partially driven by consumers’ perceptions that organic food is more nutritious (34). Motivation for buying organic chicken was perceptions that it had less residues of antibiotics and chemicals, and was safer and healthier than non-organic chicken (30). According to published literature organic food is not more nutritious than conventional foods (34,35). However, consumption of organic foods may reduce exposure to pesticide residues and antibiotic-resistant bacteria (35).

The informants in our study believed that Swedish consumers favour Swedish-produced food, and perceived Swedish chicken meat to be safer and of better quality. This factor was used in the marketing of chicken meat and ‘buying Swedish’ had a positive connotation. Despite this, there seems to be an everlasting struggle to promote Swedish products in order to keep their position on the market. All our informants worried about the threat from non-Swedish low-cost food and said that Swedish food production must be protected, for instance by educating consumers to make them aware of how Swedish food producing animals are raised. A study from Finland revealed that when Finnish consumers were told about animal welfare in production, the food production method became a more important factor for them in their choice of food (32). In Sweden there is a trend of consumers buying ecological or locally small-scale produced food, but the cost is often high. The informants presented the Swedish large-scale production of chicken-meat as a sustainable alternative, which they hope will continue to be the choice of many customers.

### Economy rules

To decrease the need of antibiotics globally new production methods must be introduced in many countries (19,20). Changing production methods means higher production costs, which has been identified as a hindering factor for reducing antibiotic use, as well as reducing capacity for reinvestment in farm buildings (15). In countries like India where regulations are in early phases, effective control strategies are lacking (18). Considerable disparities in testing practices on antibiotic usage and antibiotic resistance levels have been identified and methods must be harmonised to allow for comparability (27). Investments in preventive measures are necessary and this is a matter for policymakers and authorities (2). However, as veterinarians in our study concluded, a country with less financial resources will have challenges advancing as far as for instance Sweden has. Maybe it is time to take a global perspective and discuss cost-sharing and suggest that rich countries contribute not only with knowledge but also economical resources to countries now in the process of developing their antibiotic resistance containment measures.

### One Health approach

The One Health approach means that measures must be taken in human, animal and environmental sectors and that actions should be coordinated (2,6). Antibiotic resistance can be looked at from different perspectives and accordingly be described as different problems that need different strategies (36). Suggested perspectives are antibiotic resistance as healthcare, as development, as innovation, as security and as One Health. One Health was said to already provide a converging way to conceptualise and address antibiotic resistance (36).

In Sweden the One Health approach was implemented at policy level but not among practitioners. The containment of antibiotic resistance in Sweden primarily engages the sectors separately and efforts have been going on for a long time in both animal and human sector. For countries starting their journey towards lower antibiotic use, it can probably help with a One Health approach. Sweden has worked for 20-30 years to get where it is today. That time does not exist for countries about to start their work now. Hopefully, a strategy based on One Health will help and be more effective. Also, it will be interesting to see how the One Health approach will influence antibiotic resistance containing measurements in Sweden.

### Limitations and strengths

Trustworthiness is of major importance in all research. In this study we used the criteria developed by Lincoln and Guba to ensure high quality (37,38). To meet the criteria of credibility we recruited stakeholders with different experience to gain a broad view of perceptions. Furthermore, the analysis was well structured and carefully performed. Quotations from the text are used to demonstrate confirmability. Transferability must be judged by the readers themselves and to make this possible we described how the data were collected and analysed and gave background information about the participants. Due to practical and financial reasons the number of informants was limited. The study included a small number of Swedish stakeholders, and practitioners had experience from poultry sector. The poultry farmers were recruited via the veterinarians, and it is possible that they had more knowledge of antibiotics and were more motivated to work according to guidelines than farmers in general. However, we recruited one egg farmer separately and used this interview to get a wider picture. This informant never used antibiotics on the farm, and awareness of antibiotic resistance was low. Like the other farmers the informant primarily described the daily work to help the animals to stay as healthy as possible. A strength of our study was the choice of personal interviews, which often give richer material, instead of by telephone, which might have produced more interviews. Additionally, our findings are in line with the perceived opinion in this field in Sweden and the consistency in responses means we feel that our findings give a good picture of knowledge, attitudes and practices in this sector.

### Conclusion

Sweden has come far in the work to contain antibiotic resistance in the animal sector by practicing restrictive use of antibiotics in food animal production. This practise is based on a long tradition of cooperation among stakeholders, from policymakers to farmers, and with a primary focus on animal health and welfare. The stakeholders were proud of the Swedish food animal production, but at the same time worried about not being able to sell their goods on the international market, as this production methods were more expensive than methods using more antibiotics.

## Acknowledgments

We thank Forte in the SAMRC-Forte collaborative research programme called SAMRC/FORTE-RFA-01-2016 for funding this study. We thank the informants for participation and sharing their views.

## Author Contributions

### Conceptualization

Cecilia Stålsby Lundborg, Jaran Eriksen, Sabiha Yusuf Essack, University of KwaZulu-Natal, Durban, South Africa

### Formal analysis

Ingeborg Björkman

### Methodology

Marta Röing, Ingeborg Björkman

### Validation

Ingeborg Björkman, Marta Röing, Cecilia Stålsby Lundborg, Jaran Eriksen

### Writing – original draft

Ingeborg Björkman

### Writing – review and editing

Marta Röing, Cecilia Stålsby Lundborg, Jaran Eriksen

